# MDM2 Inhibition in Combination with Endocrine Therapy and CDK4/6 Inhibition for the Treatment of ER-Positive Breast Cancer

**DOI:** 10.1101/2020.06.09.140921

**Authors:** Neil Portman, Heloisa H. Milioli, Sarah Alexandrou, Rhiannon Coulson, Aliza Yong, Kristine J. Fernandez, Kee Ming Chia, Davendra Segara, Andrew Parker, Sue Haupt, Ygal Haupt, Wayne D. Tilley, Alex Swarbrick, C. Elizabeth Caldon, Elgene Lim

## Abstract

**Background:** Resistance to endocrine therapy is a major clinical challenge in the management of estrogen receptor (ER)-positive breast cancer. In this setting p53 is frequently wildtype and its activity may be suppressed via upregulation of its key regulator MDM2. This underlies our rationale to evaluate MDM2 inhibition as a therapeutic strategy in treatment resistant ER-positive breast cancer.

**Methods:** We used the MDM2 inhibitor NVP-CGM097 to treat *in vitro* and *in vivo* models alone and in combination with fulvestrant or palbociclib. We perform cell viability, cell cycle, apoptosis and senescence assays to evaluate antitumor effects in p53 wildtype and p53 mutant ER positive cell lines (MCF-7, ZR75-1, T-47D) and MCF-7 lines resistant to endocrine therapy and to CDK4/6 inhibition. We further assess the drug effects in patient-derived xenograft (PDX) models of endocrine-sensitive and -resistant ER positive breast cancer.

**Results:** We demonstrate that MDM2 inhibition results in cell cycle arrest and increased apoptosis in p53-wildtype *in vitro* and *in vivo* breast cancer models, leading to potent anti-tumour activity. We find that endocrine therapy or CDK4/6 inhibition synergises with MDM2 inhibition but does not further enhance apoptosis. Instead, combination treatments result in profound regulation of cell cycle-related transcriptional programmes, with synergy achieved through increased antagonism of cell cycle progression. Combination therapy pushes cell lines resistant to fulvestrant or palbociclib to become senescent and significantly reduces tumour growth in a fulvestrant resistant patient derived xenograft model.

**Conclusions:** We conclude that MDM2 inhibitors in combination with ER degraders or CDK4/6 inhibitors represent a rational strategy for treating advanced, endocrine resistant ER-positive breast cancer, operating through synergistic activation of cell cycle co-regulatory programs.

## Background

Estrogen receptor (ER) positive breast cancer accounts for ∼ 70% of breast cancer diagnoses. For many patients, the disease is treated effectively by surgery and adjuvant endocrine therapy. Unfortunately, ∼35% of patients receiving endocrine therapy relapse with resistant disease, either through inherent resistance to treatment or the emergence of acquired endocrine resistance (1, 2). Resistant disease that retains ER positivity is typically managed by treating with additional lines of endocrine therapy. Recently, the addition of Cyclin Dependent Kinase 4 and 6 (CDK4/6) inhibitors to endocrine therapy has become the new standard of care for first and second line treatment for advanced breast cancer (3). However, even with the addition of CDK4/6 inhibitors, resistance develops on average within 2-3 years and the need to identify novel targets and therapies in the resistant setting remains pressing (3, 4).

When functioning normally, p53 is activated in response to diverse genotoxic stimuli and initiates transcriptional programmes that lead to cell cycle inhibition, DNA repair and in the case of irreversible damage, apoptosis. In ER positive breast cancer and liposarcoma, the incidence of p53 mutation is relatively low (5) but there is an increased incidence of dysregulation of the major p53 regulatory proteins, MDM2 and MDM4 (6). MDM2 and MDM4 interact directly with p53 to inhibit its transcriptional activity and promote its relocation to the cytoplasm. MDM2 additionally ubiquitinates p53 to initiate its proteosomal degradation (7-10). Consequently the pharmacological reactivation of p53 function through the inhibition of MDM2 has attracted significant attention as a potential therapeutic strategy (5).

In breast cancer, although the incidence of amplification or mutation of *MDM2* is relatively low, increased abundance of MDM2 protein occurs in ∼38% of all breast cancers and is more frequent among ER-positive than in ER-negative tumours (6, 11). There is significant interaction between the MDM2/p53 axis and ER signalling. *MDM2* is a transcriptional target of ER, and MDM2 protein interacts directly with ER (12, 13). ER also regulates and interacts with p53 (14, 15) and activation of ER by either its cognate ligand or by selective ER modulators such as tamoxifen inhibits the activity of p53 (14). Simultaneous inhibition of the MDM2/p53 interaction using small molecule inhibitors, and degradation of ER via the selective estrogen receptor degrader fulvestrant can synergistically reduce proliferation of cell line models and xenografts (14, 16). Curiously, this synergy occurs without the significant induction of apoptosis (16). An unresolved question is how MDM2 inhibition synergises with endocrine therapy, and whether outcomes would be improved in combination with the new standard-of-care treatment, CDK4/6 inhibitors.

In this study, we characterised the anti-tumour effect of p53 activation via MDM2 inhibition using the small molecule inhibitor NVP-CGM097 - a dihydroisoquinolinone derivative currently being evaluated in a phase I clinical trial (17, 18) - in endocrine-resistant and endocrine-sensitive *in vitro* and *in vivo* models of ER-positive breast cancer. We show synergistic tumour cell inhibition *in vitro* in combination with either fulvestrant or palbociclib specifically via cell cycle arrest pathways, rather than by a general upregulation of p53 activity that includes apoptosis. We then demonstrate that in endocrine- and CDK4/6 inhibitor-resistant *in vitro* models, MDM2 inhibition is potentiated by combination with endocrine therapy or CDK4/6 inhibition and that this occurs via an increase in senescence compared to MDM2 inhibition alone.

## Methods

### Cell culture and reagents

MCF7 cells (Michigan Cancer Foundation), T-47D and ZR75-1 cells (American Type Culture Collection) lines were verified through short tandem repeat profiling and tested negative for mycoplasma contamination at the commencement of the project. Frozen stocks of each line used were generated at the commencement of the project and defrosted cells were cultured for between 10 and 20 passages. Parental cell lines were cultured in RPMI 1640 media (Thermo Fisher) supplemented with 10% fetal bovine serum (GE Healthcare) and 20 mM HEPES (Thermo Fisher). Fulvestrant resistant (FasR) and palbociclib resistant (PalbR). MCF-7 derivatives were developed and cultured in phenol red-free RPMI 1640 (Thermo Fisher) supplemented with 5% charcoal stripped fetal bovine serum (GE Healthcare), 20 mM HEPES and 10pM 17β-estradiol (Sigma). Resistance to fulvestrant was developed by the continuous addition of 100nM fulvestrant (Assay Matrix) over 12 months until the cells proliferated with a constant doubling time, and cells were subsequently maintained with 100nM fulvestrant. Resistance to palbociclib was developed by the addition of 500nM palbociclib (Selleckchem) for 8 months until the cells proliferated with a constant doubling time, and cells were subsequently maintained with 500nM palbociclib. For experiments that required a no-drug treatment arm, selection was removed for 48 hours prior to commencement of treatment. Stock solutions of each drug (10,000x) were prepared in DMSO (NVP-CGM097, fulvestrant), ethanol (17β-estradiol) or water (palbociclib). All cells were cultured under 5% CO_2_ in a humidified incubator at 37°C.

### Western blot analysis

All Western blot analyses were repeated on at least three separate occasions using independently generated lysates. Cells were lysed in RIPA buffer supplemented with Halt Protease and Phosphatase Inhibitor Cocktail (Thermo Fisher). Protein lysates were separated by electrophoresis over 4–15% mini-Protean TGX gels (Bio-Rad) and transferred to 0.45µm Immobilon-FL polyvinylidene fluoride membrane (Millipore). Proteins were detected by immunoblotting with specific primary antibodies and subsequent detection by fluorescent conjugated secondary antibodies [IRDye 680RD Donkey anti-mouse (#926-68072, LI-COR) or IRDye 800CW Donkey anti-Rabbit (#926-32213, LI-COR)]. Fluorescent signals were visualised using the Odyssey CLx Imaging System (LI-COR). Anti-p21 12D1 (2947) and anti-p53 (9282) were sourced from Cell Signalling, anti-MDM2 2A10 (ab16895) was sourced from Abcam.

### Proliferation assays

Cells were seeded into 96 well plates at a density of 3000 cells in 100µl media per well. Treatment was delivered via replacement of the media in each well with fresh media containing the indicated concentrations of drugs the following day. Mitochondrial activity (as an indication of cell viability) was assessed using the alamar blue assay (Thermo Fisher) according to the manufacturer’s instructions two days after treatment. Experiments were performed in three independent replicates. Growth inhibition was calculated using the formula 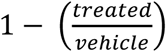. Alternatively, proliferation was measured by cell number using the IncucyteTM live-cell imaging system (Essen Bioscience). Cells were plated at 1000-3000 cells per well in a 96 well plate in 40 wells/cell line, and cell number determined by imaging every 1-2 days for 2 weeks. IC50 values were calculated from non-linear regression analyses performed in GraphPad Prism (version 7.04, GraphPad Software). Synergy was quantified using a dose-effect approach with a constant dose ratio design as per Chou and Talalay (19). Combination Indices (C.I.) were calculated using CompuSyn software (ComboSyn, Inc) such that a log_2_(C.I.) < 0 indicates synergy.

### Senescence assay

Senescence was assayed by visualising β-galactosidase activity using Senescence β-galactosidase staining kit (Cell Signalling) according to the manufacturer’s instructions. Three fields of approximately 100 cells were scored for each of three independent replicates for each treatment condition.

### Flow cytometry

Flow cytometry was performed on a Becton Dickinson CantoII flow cytometer using BD FACSDIVA software, and the results were analysed using Flowjo software (Tree Star Inc.). Cells were treated with 10nM Bromo-deoxyuridine (BrdU) for two hours prior to harvest by addition of trypsin. BrdU was detected using FITC-conjugated anti-BrdU (BD Bioscience) according to the manufacturer’s protocol. Total DNA was detected by co-staining with Propidium Iodide (PI). For quantification of apoptosis, cells were assessed using the APC AnnexinV Apoptosis Detection Kit with PI (BioLegend, CA, USA; #640932) as per the manufacturer’s instructions. Between 5000 and 10000 cells were analysed per sample and each experiment was conducted in triplicate.

### Patient-derived breast cancer xenograft (PDX) models

All *in vivo* experiments, procedures and endpoints were approved by the Garvan Institute of Medical Research Animal Ethics Committee (protocol 15/25). KCC_P_3837 (HREC/13/RPA/187) was derived from an untreated grade 3, ER-positive, PR-positive, HER2-negative primary invasive ductal carcinoma. Gar15-13 (#14002, HREC/H2006/2633) was resected from a breast cancer liver metastasis of an endocrine resistant ER-positive, PR-negative, HER2-negative metastatic breast cancer (20). At surgery, 4mm^3^ sections of tumour tissue were implanted into the 4^th^ inguinal mammary gland of 6-8 week old female NOD-SCID-IL2γR-/- mice (Australian BioResources Pty Ltd) (21). For KCC_P_3837, tumour growth was supported by implantation of a silastic pellet containing 0.72mg 17β-estradiol. Tumour growth was assessed twice weekly by caliper measurement and mice were randomized to treatment arms when tumours reached 150-250mm^3^ (using the formula: *width*^2^ × *length* × 0.5) using an online randomisation tool (https://www.graphpad.com/quickcalcs/randomize1.cfm). 100mg/kg NVP-CGM097 (20mg/ml in 2% DMSO) was administered by oral gavage 5 days a week. 5mg/body fulvestrant (50mg/ml in peanut oil – Sigma P2144) was delivered by sub-cutaneous injection once per week. Mice were treated for 60 days or until tumour volume reached 1000mm^3^.

### Immunohistochemistry (IHC) and quantification

Tumour tissue was harvested and immediately fixed in 10% neutral buffered formalin at 4°C overnight before dehydration and paraffin embedding. Antibodies used for IHC were anti-ER (M7047, 1:300, Agilent) and anti-Ki67 (M7240, 1:400, Agilent). Primary antibodies were detected using biotinylated IgG secondary antibodies (Agilent, 1:400), using streptavidin-HRP (Agilent) for amplification of signal followed by the addition of 3,3’-diaminobenzidine (Sigma) substrate. Images were scanned using Leica Aperio Slide Scanner (Leica Biosystems) and analysed using QuPath software to differentiate tumour tissue from stroma and necrosis, and to quantify Ki-67 positivity in tumour tissue.

### Statistical Analysis

Statistical analyses were performed in GraphPad Prism (version 7.04, GraphPad Software) or in Microsoft Excel (χ^2^ test for cell cycle phase analysis). * = p<0.05, ** = p<0.01, *** = p<0.001, **** = p<0.0001, ns = p>=0.05.

### Differential Expression and Gene Set Enrichment Analysis

Total RNA was extracted from breast cancer cell lines and tissue samples using RNeasy Mini Kit (Qiagen) following the manufacturer’s instructions. For tumour tissue, snap frozen sections were first disrupted in lysis buffer in a TissueLyser II (QIAGEN). Sequencing data was processed on the Illumina HiSeq 2500 v4.0 System. Sequences were trimmed using Trim Galore (version 0.4.4) and genome-guided alignment to a human reference (HG38) was performed using STAR software (version 2.5). Feature counts were computed in R (version 3.4.3) using the package *Rsubread* (22). Differential gene expression analysis was conducted with *edgeR* (23, 24) and *limma* (25) packages, on a threshold of p.val < 0.05 and twofold change. The read count quantitation and normalized expression matrix can be downloaded from http://www.ncbi.nlm.nih.gov/geo (GSE140758). Gene set enrichment analysis (GSEA) and pathway annotation were attained using *cluterProfiler* (26) in R. GSEA was tested on hallmark and curated gene sets and on gene ontology (GO) terms from the Molecular Signatures Database (MSigDB), Broad Institute (http://software.broadinstitute.org). Pathway enrichment was analysed in the Kyoto Encyclopedia of Genes and Genomes (KEGG) database (https://www.genome.jp/kegg/pathway.html).

## Results

### The small molecule MDM2 inhibitor NVP-CGM097 inhibits ER-positive breast cancer cells in a p53-dependent manner

ER-positive primary breast cancer has a relatively low incidence of *TP53* mutation (∼20%) (5). To establish whether this situation is preserved in the resistant/metastatic setting, we assessed the frequency of *TP53* mutation or deletion across public cohorts in the cBio Cancer Genomics Portal (cBioPortal, http://cbioportal.org) (Fig. 1A). The incidence of *TP53* aberration in advanced ER-positive breast cancers was ∼20%, equivalent to that observed in primary tumours and relatively low compared to other cancer types (International Agency for Research on Cancer, IARC) database (http://p53.iarc.fr). In the same clinical datasets, no instances of *MDM2* mutation or deletion were detected (data not shown). Thus, the inhibition of MDM2 to increase p53^wt^ activity could be a viable strategy in both primary and metastatic ER-positive breast cancer settings.

**Fig 1.**
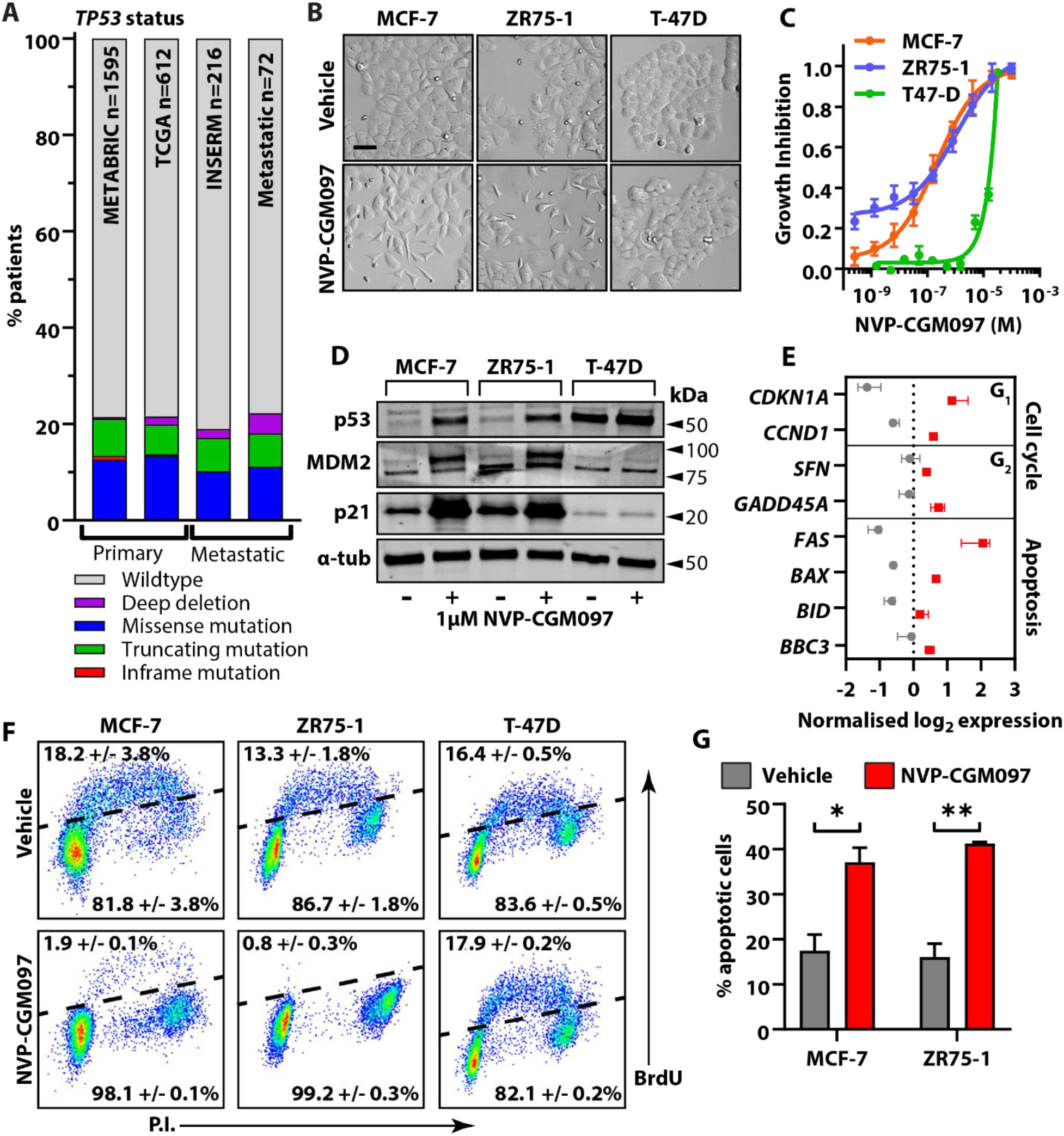
MDM2 inhibition reduces viability in a p53 dependent manner in cell line models of ER+ breast cancer. **A.** The frequency of p53 mutation (putative drivers) in ER-positive primary breast cancers – Breast Cancer (METABRIC, Nature 2012 & Nat Commun 2016), Breast Invasive Carcinoma (TCGA, Cell 2015) – and metastatic tumours – The Metastatic Breast Cancer Project (Provisional, October 2018) and Metastatic Breast Cancer (INSERM, PLoS Med 2016) – across public datasets in https://www.cbioportal.org/. **B.** Representative images of MCF-7, ZR75-1 and T-47D cell cultures after exposure to vehicle (0.01% DMSO) or 1µM NVP-CGM097 for 48 hours. Bar = 20µm. **C.** Growth inhibition relative to vehicle after 48 hours treatment with escalating doses of NVP-CGM097 in p53^wt^ MCF-7 (orange) and ZR75-1 (blue), and p53^mut^ T-47D cells (green). **D.** Western blot analysis of p53 and the known p53 transcriptional targets MDM2 and p21 after 48 hours incubation with 1µM NVP-CGM097. α-tubulin staining is shown as an indication of relative loading. See Fig. S1A for full gel and blot images. **E.** Normalised expression of cell cycle regulators and apoptosis markers after treatment with vehicle (0.01% DMSO, grey) or 1µM NVP-CGM097 (red) for 48 hours in MCF-7 cell lines. **F.** Flow cytometry analysis of the incorporation of BrdU over a 2-hour period in MCF-7, ZR75-1 and T-47D following a 48 hour incubation with 1µM NVP-CGM097. **G.** Proportion of MCF-7 and ZR75-1 cells staining positive for AnnexinV after treatment for 48 hours with vehicle (grey) or 1µM NVP-CGM097 (red). Statistical significance from Tukey’s multiple comparison test is indicated above each column.

The MDM2 inhibitor NVP-CGM097 effectively hinders cancer cell proliferation in a *TP53*^*wt*^ dependent manner (27-29). However, this has not been specifically demonstrated in ER-positive breast cancer models and we therefore determined the effect of NVP-CGM097 on cell viability in three *in vitro* models. After 48 hours of treatment, *TP53*^*wt*^ MCF-7 and *TP53*^*wt*^ ZR75-1 responded to 1µM NVP-CGM097, exhibiting a loss of normal morphology with the adoption of stellate morphology and greater separation between cells (Fig. 1B). Sub-micromolar IC_50_ concentrations were measured by alamar blue assay for MCF-7, 1.78×10^−7^M and ZR75-1, 2.05×10^−7^M (Fig. 1C). *TP53* mutant (*TP53*^*mut*^) T-47D, however, showed no morphological changes after 48 hours treatment with 1µM NVP-CGM097 (Fig. 1B) and exhibited an inhibitory response only at high doses (IC50: 7.19×10^−6^ M) (Fig. 1C).

### MDM2 inhibition activates p53 transcriptional activity in cell line models of ER-positive p53^wt^ breast cancer

Using western blot analysis, we assessed p53 reactivation in cells treated with vehicle (0.01% DMSO) or 1µM NVP-CGM097 for 48 hours (Fig. 1D, Fig. S1A). In MCF-7 and ZR75-1, p53 levels increased in response to treatment, consistent with successful inhibition of MDM2-mediated degradation of p53 protein. The p53^mut^ produced by T-47D accumulated in both vehicle and treated cells. Protein levels of the p53 transcriptional targets MDM2 and p21 were increased in MCF-7 and ZR75-1, but not in T-47D cell lines, after treatment. Collectively, these results indicate that NVP-CGM097 acts on-target to inhibit MDM2/p53 interaction with consequent p53^wt^ stabilisation and activation in models of ER-positive breast cancer.

We next performed RNA Sequencing (RNA Seq) on MCF-7 cell to interrogate effects on key p53 targets that are associated with cell cycle progression and programmed cell death (Fig. 1E). Treatment of MCF-7 cells with 1µM NVP-CGM097 for 48 hours induced the expression of *CDKN1A* (p21), *CCND1* (cyclin D1), *SFN* (14-3-3-σ) and *GADD45A* (Gadd45), which supports the activation of molecular mechanisms involved in cell cycle arrest. MDM2 inhibition also led to increased mRNA levels of apoptosis markers, including *FAS, BAX* (Bax), *BID* (Bid) and *BBC3* (PUMA). These findings confirm NVP-CGM097-mediated p53 activation of downstream targets at the molecular level in MCF-7 cell lines.

### MDM2 inhibition causes G_1_ and G_2_/M cell cycle arrest and apoptosis in treatment naïve breast cancer models

Given the several anti-proliferative outcomes of p53 activation, we next examined the mechanisms by which MDM2 inhibition compromises cell viability. Effects on cell cycle progression were assessed in MCF-7, ZR75-1 and T-47D cells after 48 hours in the presence or absence of 1µM NVP-CGM097. We quantified total DNA content of individual cells and the incorporation of Bromo-deoxyuridine (BrdU) over a two-hour pulse (Fig. 1F). In both MCF-7 and ZR75-1, treatment with NVP-CGM097 almost completely eliminated BrdU incorporation (χ^2^ p<0.001) with concomitant decrease in the proportion of cells in S phase and an increase in the proportion of cells in G_2_/M (MCF-7 χ^2^ p = 0.003; ZR75-1 χ^2^ p < 0.001) (Fig. S1B). In T-47D cells, there was no significant effect on either the incorporation of BrdU (χ^2^ p=0.70) or on the distribution of cells in the cell cycle (χ^2^ p=0.78) after treatment with NVP-CGM097. We quantified the effect of NVP-CGM097 on apoptosis in MCF-7, ZR75-1 and T-47D by staining for phosphatidylserine on the cell surface using fluorescent AnnexinV. Treatment with 1µM NVP-CGM097 for 96 hours significantly increased the proportion of apoptotic cells compared to vehicle treated populations in both MCF-7 and ZR75-1 (MCF-7 p<0.01; ZR75-1 p<0.001) (Fig. 1G).

The effects of MDM2 inhibition were further validated on a p53^wt^ patient derived xenograft (PDX) model of endocrine-sensitive ER-positive breast cancer derived from a surgically resected primary treatment-naïve ER-positive breast cancer (KCC_P_3837). Tumours were implanted concurrently with estradiol pellets and once tumour volume reached 100-150mm^3^ were treated with vehicle (estradiol, 2% DMSO) or NVP-CGM097 (100mg/kg by daily gavage) for 60 days. NVP-CGM097 significantly inhibited tumour growth (Fig. 2A) and final tumour volume (Fig. S1C) relative to vehicle. Immunohistological quantification of the proliferation marker Ki-67 at endpoint showed a significant decrease in the proportion of Ki-67 positive cells in tumours treated with NVP-CGM097 (Fig. 2B, Fig. S1D). Together, these data show that NVP-CGM097 is an effective inhibitor of proliferation in breast cancer models via p53-dependent induction of apoptosis and cell cycle arrest.

**Fig 2.**
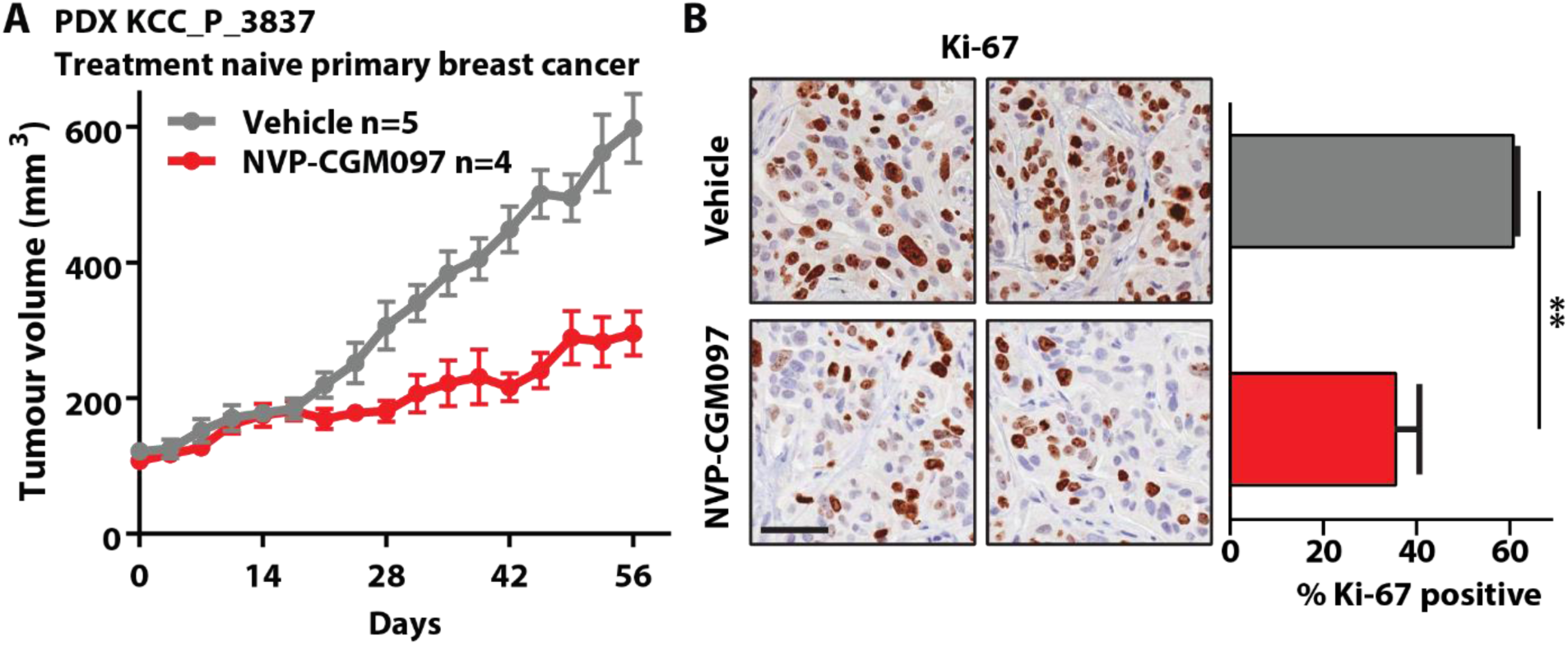
MDM2 inhibition reduces tumour growth and proliferation markers in an ER+ breast cancer PDX model. **A.** Growth of the treatment naïve KCC_P_3837 PDX model of ER+ breast cancer treated with vehicle (2% DMSO daily, grey, n=5) and NVP-CGM097 (100mg/kg daily, red, n=4) for 60 days.**B.** Representative images and quantification of the proliferation marker Ki-67 at endpoint. Ki-67 positive proportions from three independent replicates per treatment were compared by two-tailed T test. Bar = 50µm.

### The combination of NVP-CGM097 and fulvestrant synergistically inhibits MCF-7 cells without increasing apoptosis

Fulvestrant is a standard therapeutic option in the treatment of women with advanced ER-positive breast cancer. We therefore combined NVP-CGM097 with fulvestrant and measured viability of parental MCF-7 cells to test for an enhanced effect through drug additivity or synergy (Fig. 3A). We calculated combination indices using Compusyn software and detected a significant synergistic effect (log_2_ combination index < 0) on viability when combining the drugs (Fig. 3B). We then quantified the effect of NVP-CGM097 alone and in combination with fulvestrant on the rate of apoptosis in MCF-7 cells (Fig. 3C). As previously shown (Fig. 1F), treatment with 1µM NVP-CGM097 for 96 hours significantly increased the proportion of apoptotic cells compared to vehicle treated populations (p<0.01). By contrast, the inclusion of 1nM fulvestrant yielded no significant additional effect, suggesting that the synergy observed in the viability assays does not occur via enhancement of p53-mediated apoptosis.

**Fig 3.**
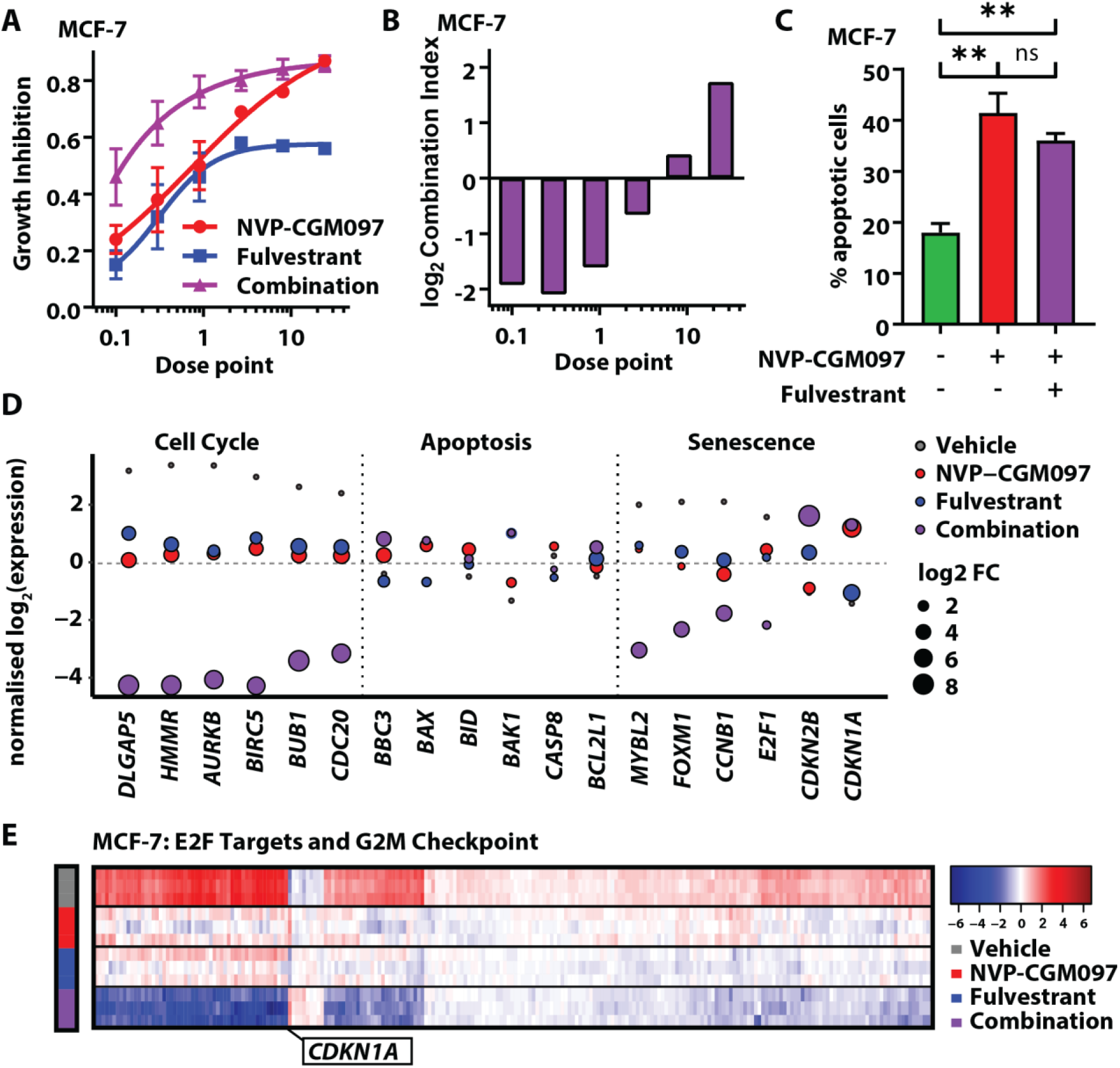
MDM2 inhibition and endocrine therapy synergistically reduce proliferation and downregulate hallmark gene sets of cell cycle progression. **A.** Growth inhibition relative to vehicle of escalating doses of NVP-CGM097 (red) and fulvestrant (blue) alone and in combination (purple) **B.** log_2_ (combination index) calculated from results in **A.** Values below 0 (equivalent to combination index < 1) indicate synergy. Dose points: NVP-CGM097 x100nM; fulvestrant x0.1nM. **C.** Proportion of MCF-7 cells staining positive for Annexin V (apoptotic cells) after treatment for 48 hours with vehicle (green), 1µM NVP-CGM097 (red), or the combination of 1µM NVP-CGM097 and 1nM fulvestrant (purple). Statistical significance from Tukey’s multiple comparison test is indicated above each column**. D.** Normalised log2 expression of significant p53 target genes induced by treatment for 48 hours with NVP-CGM097 (1μM, red), fulvestrant (1nM, blue) and combination (1µM NVP-CGM097 and 1nM fulvestrant, purple), averaged by condition, associated with cell cycle, apoptosis and senescence. **E.** Heat map of significant downregulated genes from the E2F Targets and G_2_/M Checkpoint Molecular Signatures Database (MSigDB) gene sets, normalised by gene (row), in MCF-7 cell lines for each condition: vehicle (0.01% DMSO), NVP-CGM097 (1μM), fulvestrant (1nM) and combination (1µM NVP-CGM097 and 1nM fulvestrant).

### Combination of NVP-CGM097 with fulvestrant downregulates E2F and G_2_/M transcriptional programmes *in vitro*

To identify the molecular mechanisms underlying the combination effects of NVP-CGM097 and fulvestrant we quantified the transcriptional profile of MCF-7 cells after 48 hours treatment with 0.01% DMSO, 1µM NVP-CGM097, 1nM fulvestrant or the combination of NVP-CGM097 and fulvestrant at these doses (Fig. S2A). Under stringent thresholds (adjusted p-value<0.05- and 2- fold change), 576 genes were significantly regulated by NVP-CGM097 Table S1), 803 by fulvestrant (Table S2) and 1104 by combination (Table S3) compared to vehicle. Combination therapy incorporated most of NVP-CGM097 (80%) and fulvestrant (90%)-induced gene regulation, and further altered unique gene sets that are central players in DNA replication, including the minichromosome maintenance protein complex (MCM) and DNA polymerase subunits (Fig. S2B, Table S4). The involvement of these additional gene sets suggests direct and indirect downstream regulation of the cell cycle machinery.

Kyoto Encyclopedia of Genes and Genomes (KEGG) pathway enrichment analysis of differential gene expression induced by combination therapy revealed a significant negative regulation of *Cell Cycle* (p=1.08E-21; Fig. S2C) and positive enrichment of *p53 Signalling Pathway* (p=1.95E-10; Fig. S2, C and D; Table S5) compared to vehicle. NVP-CGM097 in combination with fulvestrant significantly regulated a set of genes involved in intertwined signalling cascades of cell cycle, apoptosis and cellular senescence (Fig. 3D; Fig. S3A). In MCF-7 cell lines, combination therapy enhanced the downregulation of essential components of the cell cycle (*HMMR, AURKB, BIRC5*) that control proliferative activities and tumour growth alongside downregulation of the anti-senescence markers *FOXM1* and *E2F1* (30). FOXM1 also regulates the expression of G_2_/M cell cycle genes, including the levels of its known mitotic targets *CCNB1* and *CDC20* (31, 32). The upregulation of pro-senescence markers *CDKN1A* (p21) and *CDKN2B* (p15) – master effectors of multiple tumour suppressor pathways – and pro-apoptotic markers, such as *BBC3, BAX, BID* and *BAK1*, also contributes to the regulation of cell homeostasis, although our *in vitro* data of combination therapy indicate that synergy between treatments is not dependent on increased apoptosis.

Gene Set Enrichment Analysis (GSEA) revealed enhanced downregulation of key hallmarks involved in cell proliferation and cell cycle regulation, including E2F Targets and G_2_/M Checkpoint, in combination treated cells compared to either drug alone (Fig. 3E). To quantify the additional effect elicited by combination therapy at the transcriptional level, we computed a score for each gene *x* based on the log fold change compared to vehicle for each condition (Tables S1-3) as follows: *Score*_*x*_ = *log*2(*Combination*_*x*_/(*NVPCGM*097_*x*_ + *fulvestrant*_*x*_)) (Table S6). Using this formula, a score greater than 0 denotes transcripts in which the differential gene expression detected for the combination treatment exceeds the sum of effect for each individual treatment. Transcripts from the KEGG annotated Cell Cycle gene set were significantly over-represented within the cohort scoring above 0 (Wilcoxon rank sum test p=4.45×10^−7^)

### The combination of MDM2 inhibition and fulvestrant increases senescence in fulvestrant resistant cells compared to MDM2 inhibition alone

We next asked whether combining NVP-CGM097 and fulvestrant could lead to enhancement or re-sensitisation to therapy in fulvestrant resistant MCF-7 (FasR) cell lines. MCF-7 FasR were generated by the continuous administration of 100nM Fulvestrant for 12 months to MCF-7 cells. Resistance was confirmed by examination of relative proliferation of the parental MCF-7 cells and MCF-7 FasR cells using an Incucyte confluence assay (Fig. S4A). We first treated MCF-7 FasR cells with escalating doses of NVP-CGM097 in the presence or absence of 100nM fulvestrant (the dose used to generate fulvestrant resistance in this model) (Fig. 4A). No significant difference was observed in cell viability between the two treatments using the alamar blue assay. Treatment with 1µM NVP-CGM097 for 48 hours significantly altered the cell cycle phase distribution of MCF-7 FasR cells (χ^2^ p<0.001) resulting in a decrease in the proportion of cells in S phase, consistent with our findings from parental MCF-7 cells, and no significant difference in distribution was observed with the addition of 100nM fulvestrant (Fig. 4B). Increased levels of p21 protein were comparable between treatment with 1µM NVP-CGM097 alone for 48 hours and after treatment in combination with 100nM fulvestrant (Fig. 4C, Fig. S4B). Treatment with 100nM fulvestrant alone yielded no change in p21 levels compared to untreated cells. We observed phenotypically distinct cells following treatment with NVP-CGM097 alone and in combination with fulvestrant, with a flattened morphology reminiscent of senescence (Fig. 4D). Consequently, we analysed the cells for senescence via accumulation of senescence-associated β-galactosidase. We observed a significant increase in β-galactosidase staining in cells treated with the combination of 1µM NVP-CGM097 and 100nM fulvestrant for 72 hours compared to vehicle treatment or treatment with either single agent alone (p<0.01 vs NVP-CGM097; p<0.001 vs fulvestrant) (Fig. 4E).

**Fig 4.**
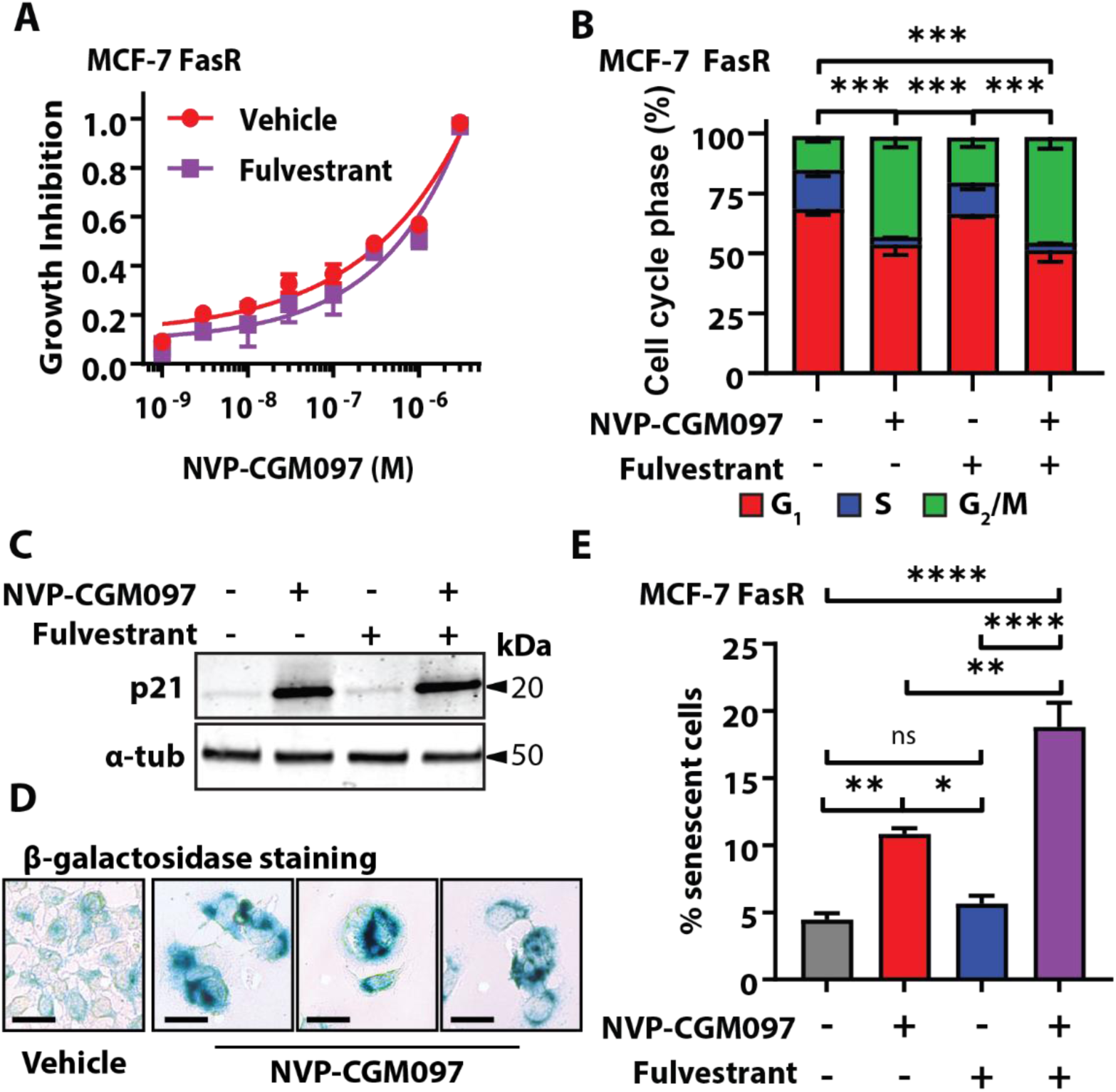
Fulvestrant potentiates MDM2 inhibition in a fulvestrant resistant cell line model. **A.** Growth inhibition of MCF-7 FasR relative to vehicle with escalating doses of NVP-CGM097 in the presence (purple) or absence (red) of 100nM fulvestrant – the dose used to generate the resistant cell line. No significant difference was observed after 48 hours of treatment. **B.** Flow cytometric quantification of propidium iodide incorporation into DNA compared to vehicle in MCF-7 FasR after incubation for 48 hours with vehicle (0.01% DMSO), 1µM NVP-CGM097, 100nM fulvestrant, or the combination of both drugs. Statistical significance from χ^2^ test using the vehicle treated profile as the expected value is indicated. Red = G_1_ (bottom), blue = S (middle), green = G_2_/M (top). **C.** Western blot analysis of the p53 transcriptional target p21 in MCF-7 FasR cells after 48 hours incubation with 1µM NVP-CGM097. α-tubulin staining is shown as an indication of relative loading. See Fig. S4B for full gel and blot images. **D.** Representative images of MCF-7 FasR cell cultures after exposure to vehicle (0.01% DMSO) or 1µM NVP-CGM097 for 48 hours followed by staining for senescence associated β-galactosidase. Bar = 20 µm **E.** Quantification of MCF-7 FasR cells staining positive for senescence associated β-galactosidase activity after treatment for 48 hours with vehicle (0.01% DMSO, grey), 1µM NVP-CGM097 (red), 100nM fulvestrant (blue), or the combination of NVP-CGM097 and fulvestrant (purple). Statistical significance from Tukey’s multiple comparison test is indicated above each column.

### Combination NVP-CGM097/fulvestrant reduces tumour growth in an *in vivo* PDX model of endocrine resistance

To investigate this enhancement of effect *in vivo*, we treated a p53^wt^ PDX model of endocrine-resistant ER-positive breast cancer, Gar15-13 (20), with vehicle (DMSO), NVP-CGM097, fulvestrant or combination therapy. Gar15-13 was established from a liver metastasis progressing on the aromatase inhibitor anastrozole. It maintains high ER positivity but grows independently of estradiol supplementation, consistent with its prior exposure to aromatase inhibition in the patient (20). This model is resistant to fulvestrant and fulvestrant-treated tumours progressed rapidly to tumour volume endpoint with growth kinetics and survival outcomes similar to the vehicle arm (Fig. 5A). NVP-CGM097 alone significantly inhibited tumour growth (Fig. 5A) and extended survival (Fig. 5, B and C) and surprisingly, given the model’s resistance to fulvestrant, combination treatment reduced tumour growth (Fig, 5A) and increased survival time significantly more than NVP-CGM097 alone, with benefit continuing beyond the six week period of administration (Fig. 5, B and C). Gar15-13 PDX treated for ten days with NVP-CGM097 alone or in combination with fulvestrant showed reduced Ki-67 staining compared to vehicle treated tumours, although no significant difference was observed between the NVP-CGM097 only arm and the combination arm (Vehicle vs NVP-CGM097 p<0.01; Vehicle vs combination p=0.011, all other pairwise comparisons not significant at α=0.05) (Fig. 5D). Tumours treated for ten days with fulvestrant (either alone or in combination) showed a significant reduction in the intensity of ER staining by immunohistochemistry in both the nucleus and cytoplasm (Vehicle vs fulvestrant -nuclear ER p=0.038, cytoplasmic ER p<0.01; vehicle vs combination - nuclear ER p=0.013, cytoplasmic ER p<0.001) (Fig. S4C), demonstrating efficacy of fulvestrant in targeting ER but not in directly inhibiting growth. The observed enhancement of the effects of NVP-CGM097 with the addition of fulvestrant may therefore be the result of an interaction with functions of ER not directly associated with proliferation, for example, in the context of senescence as we have demonstrated in MCF-7 FasR.

**Fig 5.**
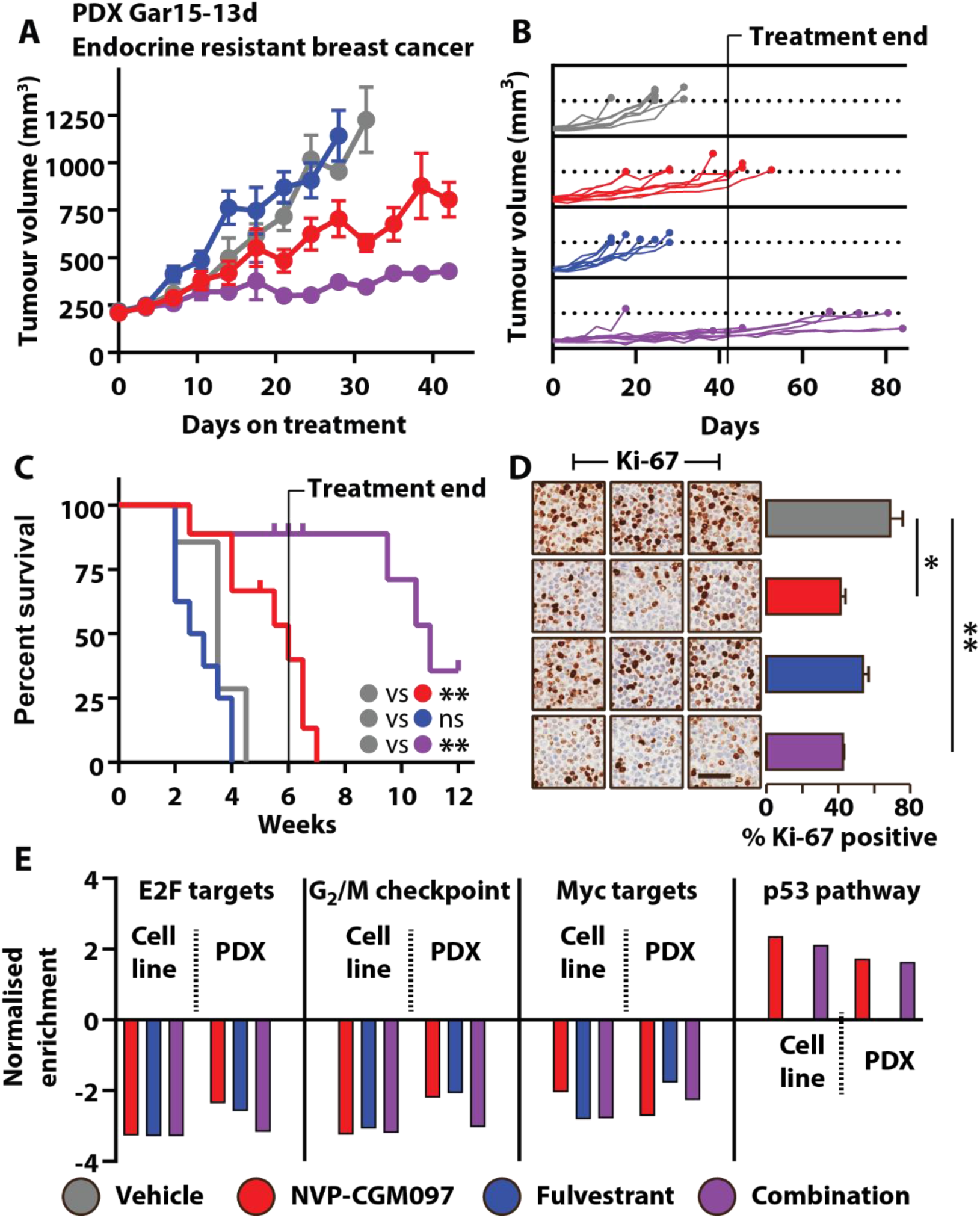
Fulvestrant potentiates MDM2 inhibition *in vivo* in a PDX model of fulvestrant resistant ER+ breast cancer. **A.** Growth of the endocrine resistant Gar15-13 PDX model of ER+ breast cancer treated with vehicle (2% DMSO daily, grey, n=7), NVP-CGM097 (100mg/kg daily, red, n=8), fulvestrant (5mg/body weekly, blue, n=8) or the combination of NVP-CGM097 and fulvestrant (purple, n=9).**B.** Continued growth of tumours following withdrawal of treatment after 6 weeks. **C.** Survival analysis of outcomes after treatment with combination therapy compared to each drug alone (an event was defined as a tumour exceeding 999 mm^3^). Curves were compared using Mantel-Cox log-rank test and significance is indicated for comparisons of each arm with vehicle. **D.** Representative images and quantification of IHC staining for Ki-67 of three independent replicates per treatment arm from tumours treated for 10 days. Statistical significance from Tukey’s multiple comparison test is indicated. Bar = 50µm. **E.** GSEA analysis compared to vehicle performed across gene expression sets of *in vitro* and *in vivo* models following treatment with NVP-CGM097 (red), fulvestrant and combination therapy show consistent regulation of hallmark signatures from MSigDB: E2F Targets, G_2_/M Checkpoints, MYC Targets and p53 Pathway. This analysis is reinforced by the normalized enrichment score (NES) that shows the difference in gene set size following each treatment conditions.

### Effects of single agent and combination therapy reconcile targeted molecular mechanisms in both *in vitro* and *in vivo* models

RNASeq analysis of short-term (10 day) treated tumours showed differential genes (p<0.05) induced by NVP-CGM097 (n=141), fulvestrant (n=411) and combination therapy (n=484) (Table S7-9). GSEA demonstrated negative enrichment of E2F Targets and G_2/_M Checkpoint and other relevant hallmark gene sets, including MYC Targets and p53 Pathway (Table S10-15). These signatures were concurrently enriched in both *in vitro* and *in vivo* models (Fig. 5E). In Gar15-13, combination therapy elicited an enhanced effect on cell cycle proliferative gene sets, E2F and G_2_/M (Fig. S4D and E; Table S16) compared to single agent treatments. Notably, fulvestrant treatment *in vitro* or *in vivo* did not show effect on the hallmarks of p53 Pathway (Fig. 5E), which was reassuringly strongly upregulated by treatment with NVP-CGM097 in both settings.

Combination treatment of NVP-CGM097 with fulvestrant also led to decreased expression of key cell cycle markers, including *DLGAP5, HMMR, AURKB, BIRC5, BUB1* and *CDC20* (Fig. S4E), which is consistent with those identified in MCF-7 treated cells. The strong inhibitory effect of combination therapy on cell cycle thus provide a preclinical rationale to use MDM2 inhibitors with ER degraders to enhance the therapeutic effects of endocrine sensitive and resistant ER-positive p53^wt^ breast cancer.

### MDM2 inhibition synergises with CDK4/6 inhibitors via the same mechanisms that underlie synergy with fulvestrant

Treatment with CDK4/6 inhibitors has become a standard of care for advanced ER positive breast cancer that has failed endocrine therapy (4). CDK4/6 inhibitors act through the G_1_/S axis to prevent cell cycle entry and have many overlapping target genes with endocrine therapy. Consequently, we investigated the potential of combination therapy of NVP-CGM097 and palbociclib, a major CDK4/6 inhibitor used in clinical practice. We performed a constant dose ratio protocol with NVP-CGM097 and palbociclib (Fig. 6A), followed by combination index analysis on MCF-7 cells. A significant synergistic effect on viability was observed with the combination therapy compared to either single agent alone (Fig. 6B).

**Figure 6.**
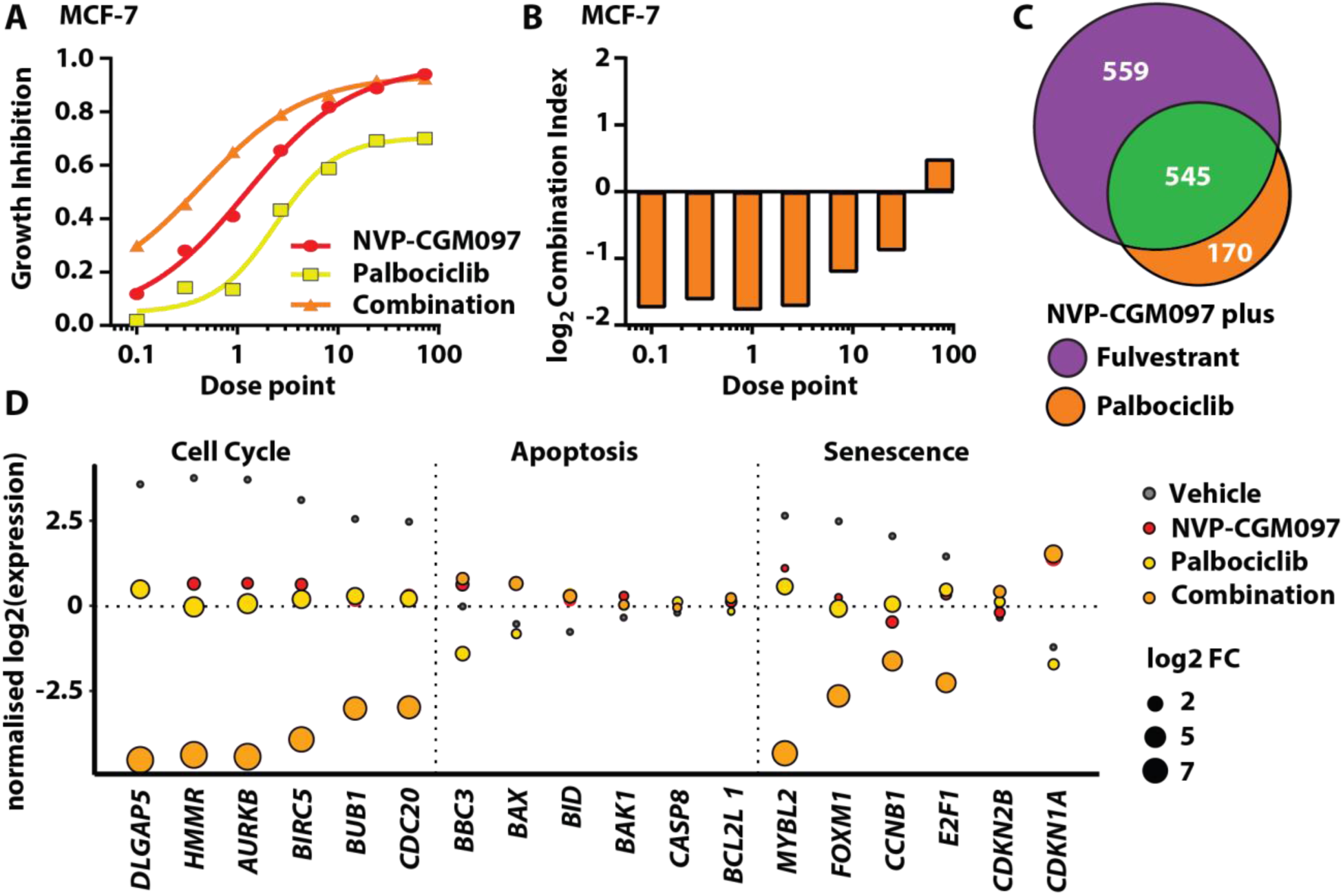
MDM2 inhibition synergises with palbociclib to reduce proliferation via cell cycle arrest. **A.** Growth inhibition relative to vehicle of escalating doses of NVP-CGM097 (red) and palbociclib (yellow) alone and in combination (orange) **B.** log_2_ (combination index) calculated from results in **A.** Values below 0 (equivalent to combination index < 1) indicate synergy. Dose points: NVP-CGM097 x100nM; palbociclib x50nM. **C.** Venn diagram showing the overlap (green) between differentially expressed genes (adjusted p-value < 0.05- and 2-fold change) induced in MCF-7 cells by treatment for 48 hours with combination therapies, 1µM NVP-CGM097 plus 1nM fulvestrant (purple) and 1µM NVP-CGM097 plus 500nM palbociclib (orange). **D.** Normalised log_2_ expression levels of significant p53 target genes induced by treatment with NVP-CGM097 (1µM); palbociclib (500nM) and combination (1µM NVP-CGM097 and 500nM palbociclib), averaged by condition, associated with cell cycle, apoptosis and senescence.

We next investigated the combinatorial effect of NVP-CGM097 and palbociclib by RNA-Seq (Fig. S5A). We confirmed significant (adjusted p-value<0.05- and 2-fold change) gene expression changes induced by NVP-CGM097 (n=572), palbociclib (n=392) and combination therapy (n=714) (S17-19), compared to vehicle (0.01% DMSO). Combination therapy encompassed about 90% of the changes promoted by either agent alone (Fig. S5B). Combination of NVP-CGM097 with fulvestrant or with palbociclib showed a significant overlap (Fig. 6C). As with NVP-CGM097 plus fulvestrant, when we quantified the additional effect elicited by combination therapy at the transcriptional level (Table S20), transcripts from the KEGG annotated Cell Cycle gene set were significantly over-represented within the cohort scoring above 0 (Wilcoxon rank sum test p=1.52×10^−13^). Combination of NVP-CGM097 with palbociclib further downregulates the senescence gatekeepers *MYBL2, FOXM1, CCNB1* and *E2F1* (Fig. 6D). Although both fulvestrant and palbociclib interact with the p53 pathway at numerous points, especially fulvestrant which ablates the interaction between ER and p53 (14), our data suggest that the major common mechanistic events underlying the efficacy of the two combination strategies with NVP-CGM097 are related to the activity of each drug in terms of inhibiting cell cycle progression.

### The combination of MDM2 inhibition and palbociclib increases senescence in palbociclib resistant cells compared to MDM2 inhibition alone

We investigated the effect of MDM2 inhibition on MCF-7 cells with acquired resistance to palbociclib (PalbR). MCF-7 PalbR were generated by the continuous administration of 500nM palbociclib for 6 months to MCF-7 cells. Resistance was confirmed by examination of relative growth inhibition of the parental MCF-7 cells and MCF-7 PalbR cells to a range of doses of palbociclib (Fig. S5C). NVP-CGM097 inhibited growth of MCF-7 PalbR cells with an IC_50_ of 9.0×10^−8^M (95% C.I. 6.7×10^−8^M to 1.1×10^−7^M), as determined by alamar blue assay (Fig. 7A). Analysis of cell cycle phase by flow cytometry showed a significant difference (χ^2^ p<0.001) in distribution from vehicle (500nM palbociclib, 0.01% DMSO) following treatment for 48 hours with 1µM NVP-CGM097 (Fig. 7B) with an increase in the proportion of cells in G_2_/M. A key difference to endocrine resistance models was that MCF-7 PalbR cells did not accumulate in G_1_ after NVP-CGM097 treatment. This suggests that cells resistant to CDK4/6 inhibition may be able to overcome the G_1_ block associated with activation of p53 to some extent but are still susceptible to cell cycle inhibition in S and/or G_2_/M. Treatment with 1µM NVP-CGM097 for 96 hours resulted in a significant increase in the proportion of apoptotic cells compared to vehicle (0.01% DMSO) or 500nM palbociclib treated cells (p<0.001) but the combination treatment yielded no significant increase compared to NVP-CGM097 alone (Fig. 7C).

**Figure 7.**
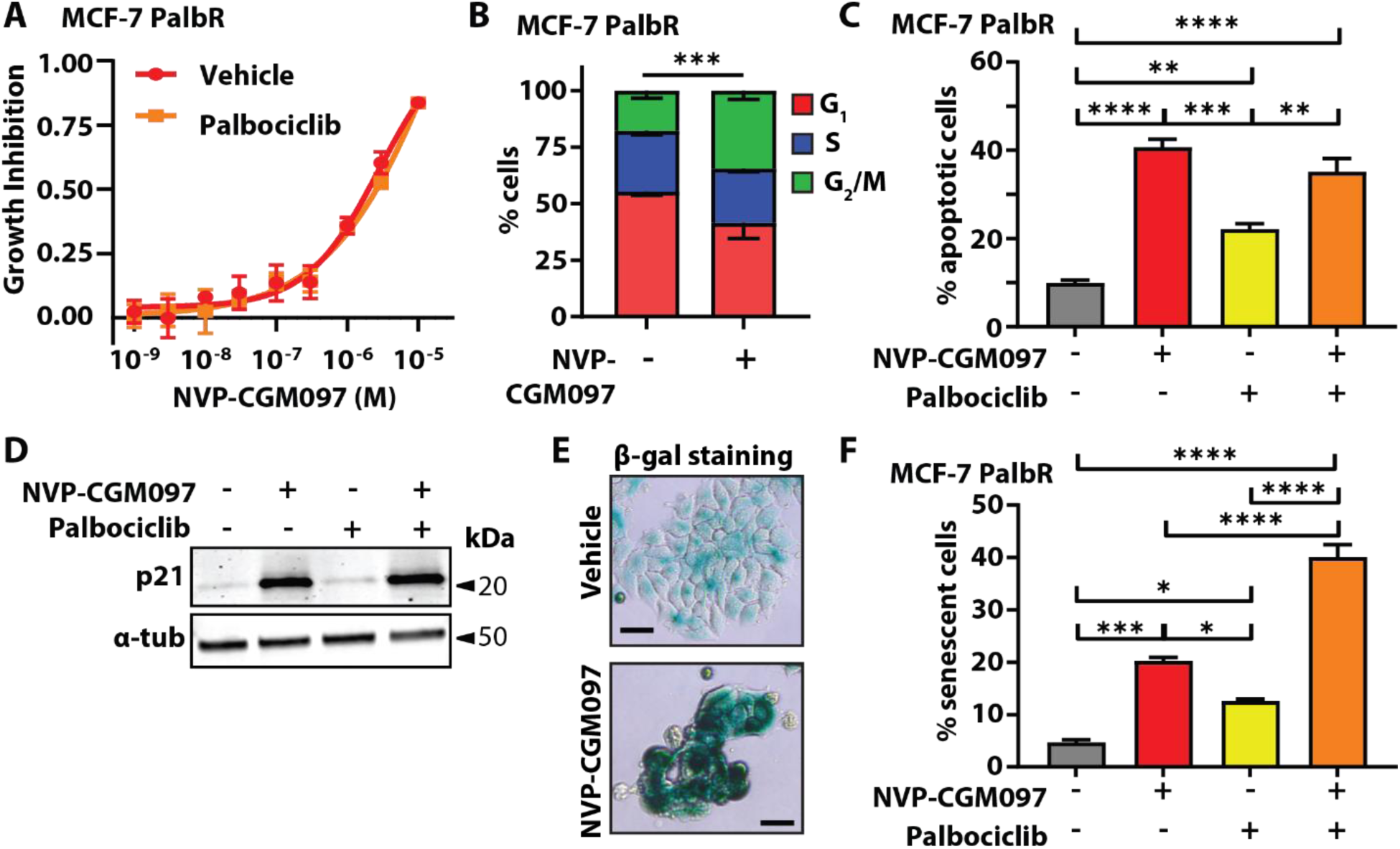
Palbociclib potentiates MDM2 inhibition via an increase in senescence in palbociclib resistant MCF-7. **A.** Growth inhibition of MCF-7 PalbR relative to vehicle (0.01% DMSO) of escalating doses of NVP-CGM097 in the presence (red) or absence (yellow) of 500nM palbociclib – the dose used to generate the resistant cell line. No significant difference was observed after 48 hours of treatment. **B.** Proportion of cells in each cell cycle phase from flow cytometric quantification of propidium iodide staining of genomic DNA in MCF-7 PalbR cells after 48 hours treatment with 1µM NVP-CGM097 compared to vehicle (500nM palbociclib, 0.01% DMSO). Statistical significance from χ^2^ test using the vehicle treated profile as the expected value is indicated. Red = G_1_ (bottom), blue = S (middle), green = G_2_/M (top). **C.** Proportion of MCF-7 PalbR cells staining positive for AnnexinV after treatment for 48 hours with vehicle (0.01% DMSO, grey), 1µM NVP-CGM097 (red), 500nM palbociclib (yellow) or the combination of 1µM NVP-CGM097 and 500nM palbociclib (orange). Statistical significance from Tukey’s multiple comparison test is indicated above each column. **D.** Western blot analysis of p21 in MCF-7 PalbR cells after 48 hours incubation with 1µM NVP-CGM097. α-tubulin staining is shown as an indication of relative loading. See Fig. S5D for full gel and blot images. **E.** Representative images of MCF-7 PalbR cell cultures after exposure to vehicle (500nM palbociclib, 0.01% DMSO) or 1µM NVP-CGM097 for 48 hours followed by staining for senescence associated β-galactosidase. Bar = 20µm **F.** Quantification of MCF-7 PalbR cells staining positive for senescence associated β-galactosidase activity after treatment for 48 hours with vehicle (0.01% DMSO, grey), 1µM NVP-CGM097 (red), 500nM palbociclib (yellow), or the combination of NVP-CGM097 and palbociclib (orange). Statistical significance from Tukey’s multiple comparison test is indicated above each column.

We had observed that NVP-CGM097 in combination with fulvestrant resulted in increased senescence compared to NVP-CGM097 alone in fulvestrant resistant cells. We examined whether a similar increase could also be observed with CDK4/6 inhibitor-resistant cells treated with combination NVP-CGM097 and palbociclib. MCF-7 PalbR cells were treated with 1µM NVP-CGM097 in combination with 500nM palbociclib (the resistance dose) for 48 hours. Treatment with either single agent NVP-CGM097 or the combination with palbociclib increased expression of the senescence marker p21 (33) (Fig. 7D, Fig S5D) and there was a significant increase in the proportion of β-galactosidase positive cells in the combination treatment compared to either single agent alone (p<0.001 vs NVP-CGM097 or palbociclib) (Fig. 7, E and F). Thus, NVP-CGM097 is an effective inhibitor of ER-positive breast cancer even in the context of palbociclib resistance, and as with endocrine therapy, rechallenging cells in the presence of NVP-CGM097 increases the incidence of senescence compared with single agent treatments.

## Discussion

The treatment of patients with endocrine-resistant ER-positive breast cancer is extremely challenging, as in this setting, disease leads inevitably to death. Numerous mechanisms of treatment resistance have been identified. While some of these pathways are targetable, for instance, the PI3K pathway and the cyclinD-CDK4/6-Rb axis; the optimal next-line treatment strategies are unclear. In this context, MDM2 inhibition to reactivate p53 is particularly attractive in ER-positive breast cancer as it has a relatively low incidence of *Tp53* mutation in the primary and metastatic setting (∼20%) (5). In addition, those ER-positive breast cancer patients who respond to CDK4/6 inhibitors are more likely to have intact p53 (34), meaning that there is the potential to improve the response to CDK4/6 inhibitors through p53 activation.

A number of small molecule MDM2 inhibitors have been designed and are at various stages of clinical development (35). NVP-CGM097 was one of the first of the new generation inhibitors and has advanced to phase 1 clinical trial (NCT01760525). Although the final outcomes of the trial have not yet been reported, early results showed clinical activity – predominantly stable disease – and establishment of a tolerable dose regimen, with one quarter of patients receiving treatment for at least 24 weeks (18). As with most MDM2 inhibitors that have so far reached clinical trials, side effects of NVP-CGM097 include thrombocytopenia and neutropenia which would need to be considered in any clinical application in combination with drugs with overlapping toxicity profiles such as CDK4/6 inhibitors (37). Importantly, our data show that combination of NVP-CGM097 with fulvestrant, which is not associated with hematological adverse effects, has a highly similar response profile to combination with CDK4/6 inhibition in terms of the key molecular pathways affected. It is currently unclear as to whether NVP-CGM097 itself will continue in development, however a range of next generation MDM2 inhibitors with improved tolerability are in development (38). Our results demonstrate that MDM2 inhibition is a rational target for novel therapeutic strategies in the setting of treatment resistant breast cancer.

In this study we show that reactivation of wildtype p53 using NVP-CGM097 is an effective inhibitor of tumour growth in multiple models of endocrine-naïve, endocrine-resistant and CDK4/6 inhibitor-resistant ER-positive breast cancer *in vitro* and *in vivo*. We identify the key mechanisms of the synergistic interactions between NVP-CGM097 and endocrine therapy, which occurs through the inhibition of E2F targets and G_2_/M checkpoint signalling and induction of senescence; rather than via a general upregulation of p53 dependent apoptotic pathways above the level of MDM2 inhibition alone. Moreover, we find these same pathways are specifically targeted in response to combined inhibition of both MDM2 and the CDK4/6 pathway.

Consistent with activation of p53, NVP-CGM097 induced apoptosis, and cell cycle arrest in G_1_ and G_2_, in *TP53*^*wt*^ ER-positive breast cancer cell line (MCF-7 and ZR75-1) models. The only setting in which we did not observe anti-tumour effect was in the case of the *TP53*^*mut*^ T-47D cell line model which supports our conclusion that activity of NVP-CGM097 requires wildtype p53. MDM2 inhibition was characterised by very high expression of the key cell cycle regulator p21 – which is a p53 target and principal component of cell cycle arrest in G_1_ and G_2_ and in the induction and maintenance of senescence (39).

Our data from ER positive PDX models shows that NVP-CGM097 is effective as a single agent, and it combines with fulvestrant to be highly effective in the fulvestrant-resistant Gar15-13 PDX model. In the only other breast cancer PDX experiments so far reported, another MDM2 inhibitor MI-77301 was effective at reducing tumour growth of two fulvestrant-resistant PDX models (16). Similar to our findings, co-administration of fulvestrant led to prolonged therapeutic benefit after treatment cessation in one model, although unlike our model no additional benefit was observed during treatment. The second PDX model responded so effectively to MI-77301 that an additional benefit from combination therapy could not be detected. Our investigations suggest a mechanism of action underlying the combination effect in our models and most probably in prior models. NVP-CGM097 synergised with fulvestrant and the major effect of combination was enhanced downregulation of cell cycle related transcriptional programmes, particularly the E2F regulated transcriptome that when unimpeded is activated as cells progress out of G_1_. Our data suggest that combining MDM2 inhibition with endocrine therapy leads to a profound arrest of cell cycle processes and our *in vitro* analysis of the MCF-7 FasR cell line supports this finding, consistent with an increase in senescence as a possible mechanism.

CDK4/6 inhibitors in combination with endocrine therapy are the standard of care for advanced breast cancer in the first- and second-line treatment settings. However, resistance to combination therapy will inevitably develop (4), and it is imperative that therapies that can operate in the post resistance setting are identified. We show for the first time that activation of p53^wt^ is effective in a model of CDK4/6 inhibitor-resistant breast cancer. Interestingly, we have demonstrated that p53-induced cell cycle inhibition in palbociclib-resistant MCF-7 is predominantly in the G_2_ phase. CDK4/6 inhibition acts to prevent G_1_ cell cycle entry by preventing the phosphorylation of Rb (40-42), and many mechanisms of CDK4/6 inhibitor resistance are via disruption of the G_1_/S checkpoint (4). Our findings therefore indicate that palbociclib-resistant cell lines that have evaded G_1_ checkpoints are still susceptible to G_2_ arrest. The combination of MDM2 inhibition and CDK4/6 inhibition engage pathways that remain intact in palbociclib and endocrine resistant cells, unlike the G_1_/S checkpoint which is frequently disabled in these resistance contexts.

We also show that rechallenging palbociclib-resistant cells with palbociclib in combination with MDM2 inhibition increases B-galactosidase staining and p21 expression, both indicative of senescence. Likewise, fulvestrant-resistant cells treated with combination MDM2 inhibition and fulvestrant become senescent. Senescence entry occurs from either G_1_ phase or G_2_/M phase (43). Thus, in our models of resistance to either endocrine therapy or palbociclib, NVP-CGM097-mediated induction of G_2_/M accumulation along with the downregulation of *FOXM1* and other senescence-related transcripts appears to be sufficient to trigger entry into senescence. CDK4/6 inhibitors are known to increase senescence entry and MDM2 can counteract this (44).

## Conclusions

Collectively, our results suggest that although ER has been shown to be a negative regulator of p53 (14, 45, 46), it is the anti-proliferative actions of fulvestrant (particularly where they overlap with CDK4/6 inhibition) and MDM2 inhibition interacting via cell cycle progression pathways that combine to produce an enhanced anti-tumour effect. Our data provide a preclinical rationale for the combination of MDM2 inhibition with either endocrine therapy or CDK4/6 inhibition, even in the resistant setting. Importantly, we also offer a novel mechanistic insight that will inform the future design of rational combinations with MDM2 inhibitors in ER-positive breast cancer, demonstrating the efficacy of co-targeting cell cycle pathways and perhaps suggesting an alternative strategy via designing combinations to promote the pro-apoptotic functions of p53.

## Abbreviations

ER: Estrogen Receptor
CDK4/6: Cyclin Dependent Kinase 4 and 6
C.I.: Combination Index
BrdU: Bromodeoxyuridine
P.I.: Propidium Iodide
IHC: Immunohistochemistry
GSEA: Gene Set Enrichment Analysis
GO: Gene Ontology
MSigDB: Molecular Signatures Database
KEGG: Kyoto Encyclopedia of Genes and Genomes
(T)p53wt: p53 (or TP53) wildtype
(T)p53mut: p53 (or TP53) mutant
IC50: Half maximal Inhibitory Concentration
RNASeq: RNA sequencing
PDX: Patient Derived Xenograft
FasR: Fulvestrant (Faslodex) Resistant
PalbR: Palbociclib Resistant

## Declarations

### Ethics approval and consent to participate

All *in vivo* experiments, procedures and endpoints were approved by the Garvan Institute of Medical Research Animal Ethics Committee (protocol 15/25). PDX models were collected with patient consent under approval from Royal Prince Alfred Hospital human research ethics committee (HREC/13/RPA/187 - KCC_P_3837) and St Vincent’s Hospital human research ethics committee (HREC/H2006/2633 - Gar15-13).

### Consent for publication

Not applicable

### Availability of data and materials

The datasets supporting the conclusions of this article are included within the article and its additional files. The read count quantitation and normalized expression matrix for all RNAseq experiments can be downloaded from http://www.ncbi.nlm.nih.gov/geo (GSE140758). Cell lines and PDX models not commercially available can be obtained via MTA with Garvan Institute of Medical Research.

### Authors’ contributions

NP, HHM designed and performed experiments, analysed results and wrote the manuscript. EL, CEC analysed results and wrote the manuscript. SA, RC, AY, KJF designed and performed experiments and analysed results. KMC, AS, SH, YH analysed results and provided critical feedback on the manuscript. AP provided critical reagents, oversight of IHC analyses and critical feedback on the manuscript, DS provided critical reagents, EH and WDT provided intellectual input during conception of the project and critical feedback on the manuscript.

## Acknowledgements

Not applicable

## Supplementary materials

**Table S1:** Differential expression analysis of MCF7 cell lines following 48 hours treatment with NVP-CGM097 versus vehicle (0.01% DMSO).

**Table S2:** Differential expression analysis of MCF7 cell lines following 48 hours treatment with Fulvestrant versus vehicle (0.01% DMSO).

**Table S3:** Differential expression analysis of MCF7 cell lines following 48 hours treatment with combination therapy (NVP-CGM097 + fulvestrant) versus vehicle (0.01% DMSO).

**Table S4:** Differential genes induced by combination therapy (NVP-CGM097 + fulvestrant) versus vehicle (0.01% DMSO).

**Table S5:** KEGG PATHWAY enrichment analysis of differentially expressed genes following treatment with combination of NVP-CGM097 and fulvestrant versus vehicle (0.01% DMSO).

**Table S6:** Combination effect quantified for all genes following treatment with NVP-CGM097, fulvestrant and combination across cell lines.

**Table S7:** Differential expression analysis of the PDX breast cancer model treated with NVP-CGM097 versus vehicle.

**Table S8:** Differential expression analysis of the PDX breast cancer model treated with fulvestrant versus vehicle.

**Table S9:** Differential expression analysis of the PDX breast cancer model treated with combination therapy versus vehicle.

**Table S10.** Gene set enrichment analysis of DE genes induced by NVP-CGM097 in MCF-7 cell lines.

**Table S11.** Gene set enrichment analysis of DE genes induced by fulvestrant in MCF-7 cell lines.

**Table S12.** Gene set enrichment analysis of DE genes induced by combination therapy in MCF-7 cell lines.

**Table S13.** Gene set enrichment analysis of DE genes induced by NVP-CGM097 in the PDX treated model.

**Table S14.** Gene set enrichment analysis of DE genes induced by fulvestrant in the PDX treated model.

**Table S15.** Gene set enrichment analysis of DE genes induced by combination therapy in the PDX treated model.

**Table S16:** Combination effect quantified for all genes following treatment with NVP-CGM097, fulvestrant and combination in the PDX model.

**Table S17**: Differential expression analysis of MCF7 cell lines following 48 hours treatment with NVP-CGM097 versus vehicle (0.01% DMSO).

**Table S18:** Differential expression analysis of MCF7 cell lines following 48 hours treatment with palbociclib versus vehicle (0.01% DMSO).

**Table S19:** Differential expression analysis of MCF7 cell lines following 48 hours treatment with combination therapy (NVP-CGM097 + palbociclib) versus vehicle (0.01% DMSO).

**Table S20:** Combination effect quantified for all genes following treatment with NVP-CGM097, palbociclib and combination across cell lines.

**Fig. S1.**
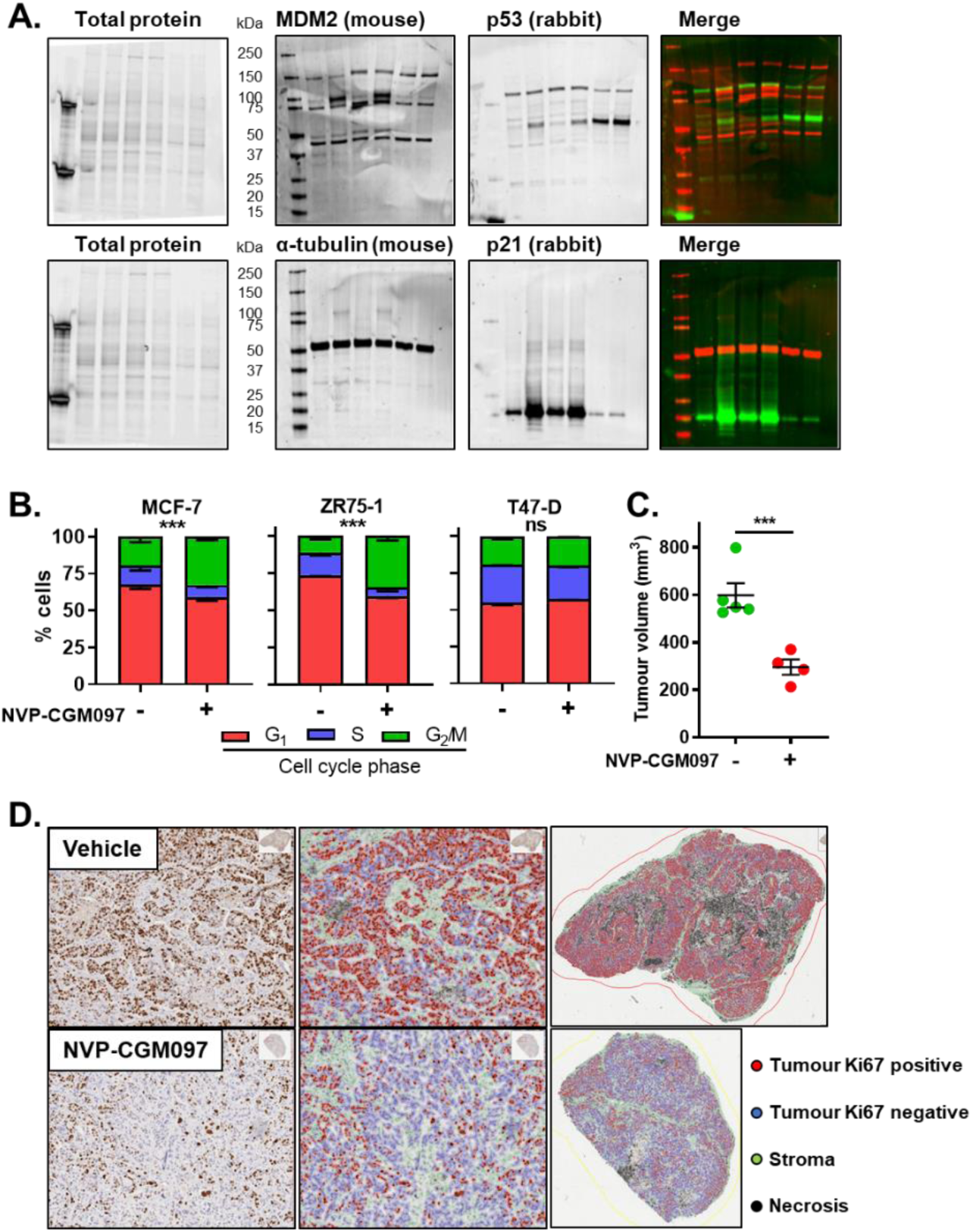
MDM2 inhibition activates p53 and reduces tumour proliferation *in vitro* and *in vivo.* **A.** Full gel and blot scans for the Western blots shown in Fig. 1B. Total protein was visualised using BioRad stain-free imaging technology according to the manufacturer’s instructions. **B.** Analysis of cell cycle phase using flow cytometry to quantify propidium iodide staining of genomic DNA shows significant alterations to cell cycle phase distribution in p53^wt^ models consistent with arrest in both G_1_ and G_2_ after incubation for 48 hours with 1µM NVP-CGM097. Red = G_1_ (bottom), blue = S (middle), green = G_2_/M (top). Statistical significance from χ^2^ test using the vehicle treated profile as the expected value is indicated. **C.** NVP-CGM097 (50mg/kg daily, red) significantly inhibited tumour volumes compared to vehicle (2% DMSO daily, green) at endpoint. Final tumour volumes were compared using two-tailed T test to determine significance. **D.** Representative images of Ki-67 quantification of endpoint tumours in Qupath software showing the classification of different tissue compartments: tumour (red and blue), stroma (green), and necrosis (black); and detection of Ki-67 negative and positive tumour cells. A single classifier was applied to all tumour sections.

**Fig. S2.**
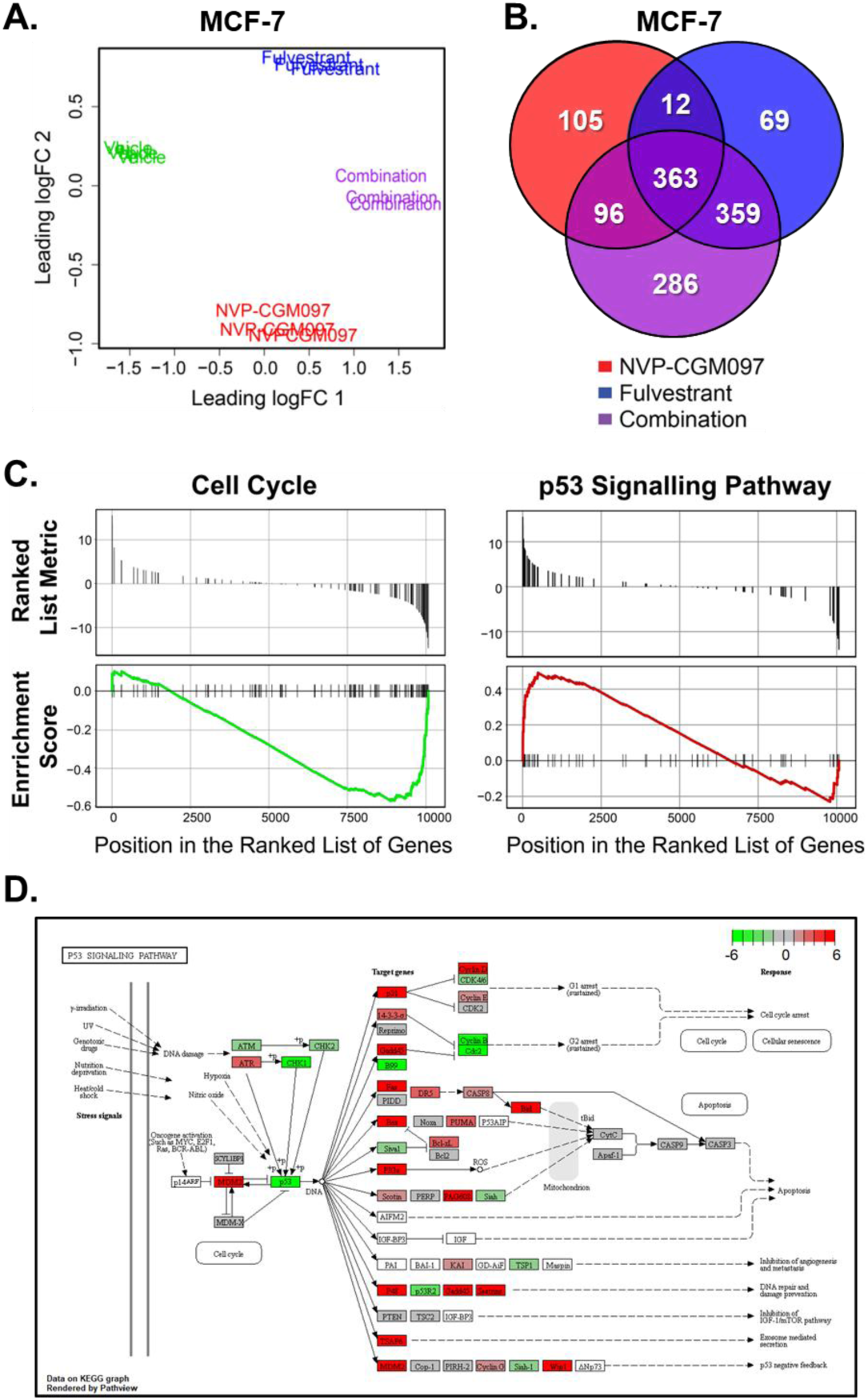
NVP-CGM097 treatment causes gene expression changes in cell cycle and p53 pathways *in vitro*. **A.** Multidimensional scaling (MDS) plot showing the level of sample similarity between MCF-7 cell lines treated with vehicle, NVP-CGM097, fulvestrant and combination therapy (NVP-CGM097 plus fulvestrant). **B.** Venn diagram showing the overlap between differentially expressed genes (adjusted p-value < 0.05- and 2-fold change) induced by treatment in MCF-7 cell lines. **C.** KEGG pathway analysis using RNA-Seq transcriptomic data shows a significant negative enrichment of Cell Cycle regulation in MCF-7 cell lines following 48 hours of treatment with 1µM NVP-CGM097 and positive enrichment of p53 Signalling Pathway. **D.** KEGG enrichment analysis showing p53 Signalling Pathway activation in MCF-7 cell lines following treatment with NVP-CGM097.

**Fig. S3.**
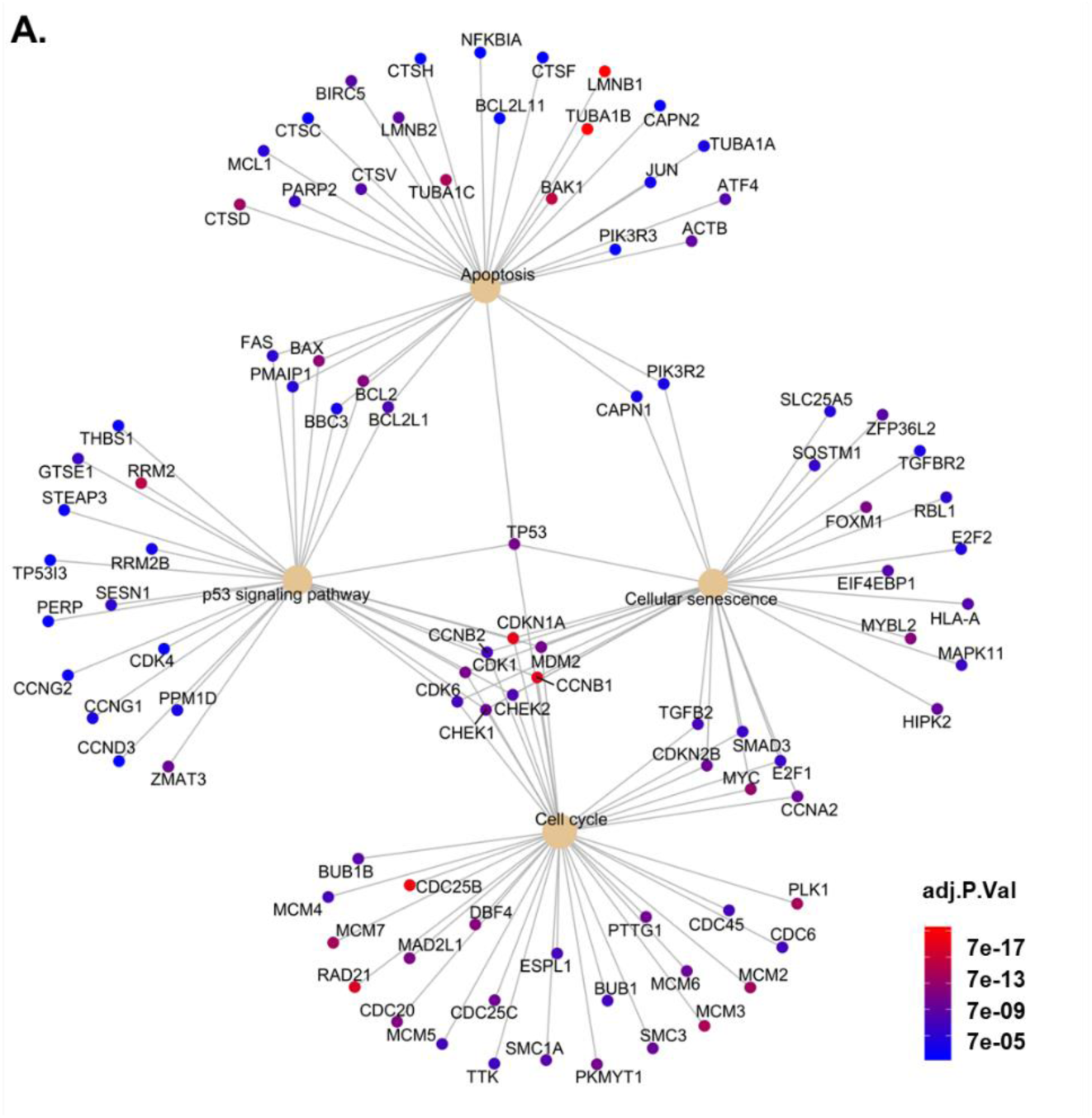
Enrichment analysis of significant differentially expressed genes against curated gene sets.

**Fig. S4.**
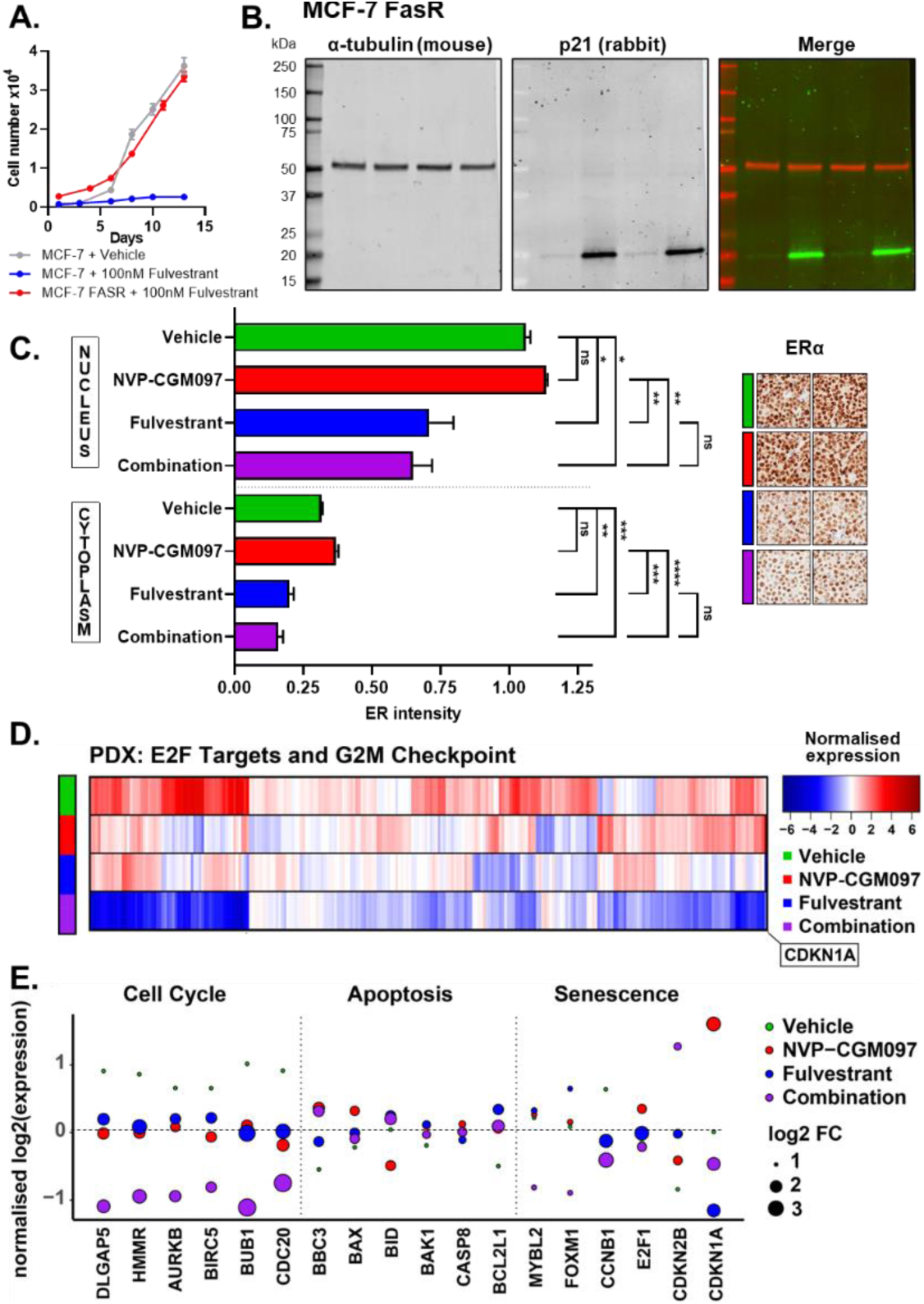
MDM2 inhibition induces changes in gene and protein expression associated with the cell cycle in endocrine resistant models *in vitro* and *in vivo*. **A.** Incucyte analysis of cell numbers of parental MCF-7 cells (blue) and MCF-7 FasR cells (red) over time after treatment with 100nM fulvestrant. Vehicle treated parental MCF-7 cell growth (grey) is shown for comparison. **B.** Full gel images for the Western blots shown in Fig. 4C. **C.** Quantification of ER staining in the nucleus and cytoplasm of three independent replicates per arm from PDX model Gar15-13 following 10 days of treatment with vehicle (2% DMSO daily, green), NVP-CGM097 (50mg/kg daily, red), fulvestrant (5mg weekly, blue) or the combination of fulvestrant and NVP-CGM097 (purple). Tumours treated with fulvestrant alone or in combination had significantly reduced ER staining in both cellular compartments compared to tumours not treated with fulvestrant. Tumour cell identification and ER quantification were performed in QuPath software. **D.** Heat map of significant downregulated genes from the E2F Targets and G_2_/M Checkpoint gene sets in the Gar1513 PDX model. **E.** Normalised log_2_ expression levels of significant p53 target genes in in the Gar1513 PDX model induced by NVP-CGM097 (red), fulvestrant (blue) and the combination of NVP-CGM097 and fulvestrant (purple), averaged by condition, associated with cell cycle, apoptosis and senescence.

**Fig. S5.**
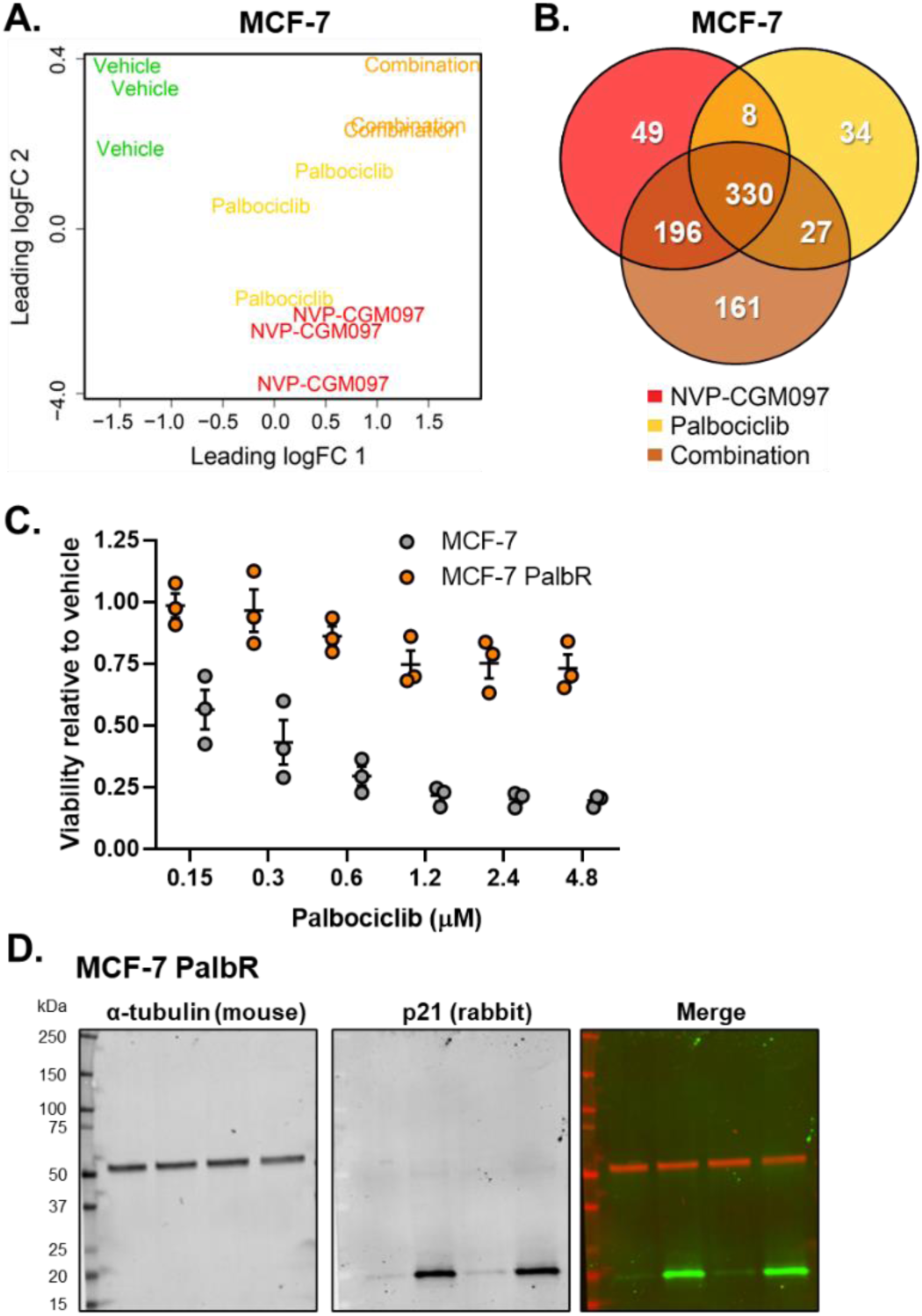
MDM2 inhibition in combination with CDK4/6 inhibition synergistically downregulates cell cycle markers *in vitro.* **A.** MDS plot of MCF-7 cell lines treated for 48h with Vehicle (0.01% DMSO), 1µM NVP-CGM097, 500nM palbociclib, or the combination of NVP-CGM097 and palbociclib. **B.** Venn diagram showing the overlap between differentially expressed genes (adjusted p-value < 0.05- and 2-fold change) induced by treatment in MCF-7 cell lines. **C.** Viability of parental MCF-7 (grey) and MCF-7 PalbR (orange) after treatment with the indicated dose of palbociclib for 72 hours. Values are shown relative to vehicle treated cells. **D.** Full gel images for the Western blots shown in Fig. 7D.

